# Ultra-high throughput single-cell RNA sequencing by combinatorial fluidic indexing

**DOI:** 10.1101/2019.12.17.879304

**Authors:** Paul Datlinger, André F Rendeiro, Thorina Boenke, Thomas Krausgruber, Daniele Barreca, Christoph Bock

## Abstract

Cell atlas projects and single-cell CRISPR screens hit the limits of current technology, as they require cost-effective profiling for millions of individual cells. To satisfy these enormous throughput requirements, we developed “single-cell combinatorial fluidic indexing” (scifi) and applied it to single-cell RNA sequencing. The resulting scifi-RNA-seq assay combines one-step combinatorial pre-indexing of single-cell transcriptomes with subsequent single-cell RNA-seq using widely available droplet microfluidics. Pre-indexing allows us to load multiple cells per droplet, which increases the throughput of droplet-based single-cell RNA-seq up to 15-fold, and it provides a straightforward way of multiplexing hundreds of samples in a single scifi-RNA-seq experiment. Compared to multi-round combinatorial indexing, scifi-RNA-seq provides an easier, faster, and more efficient workflow, thereby enabling massive-scale scRNA-seq experiments for a broad range of applications ranging from population genomics to drug screens with scRNA-seq readout. We benchmarked scifi-RNA-seq on various human and mouse cell lines, and we demonstrated its feasibility for human primary material by profiling TCR activation in T cells.

## Introduction

Microfluidic droplet generators^1-3^ are the most popular technology platform for single-cell sequencing. With their throughput, fast and simple workflow, and consistent data quality, they have contributed to the adoption of single-cell RNA-seq (scRNA-seq) in many areas of basic biology and biomedical research. In a typical droplet-based scRNA-seq experiment, the single-cell suspension is processed on a microfluidic chip, together with uniquely barcoded microbeads, reverse transcription reagents, and carrier oil (**Fig. 1a**). When aqueous and oil phases are combined at controlled flow rates, emulsion droplets co-encapsulate individual cells with individual microbeads. Cells are then lysed inside the droplets, RNA molecules anneal to bead-tethered primers, and reverse transcription is performed within the droplets. Importantly, all primers on a given bead carry the same barcode, which uniquely identifies the co-encapsulated cell. Once the transcripts have been barcoded inside droplets, the emulsion is broken and the remaining steps of the library preparation are performed in bulk.

**Figure 1:**
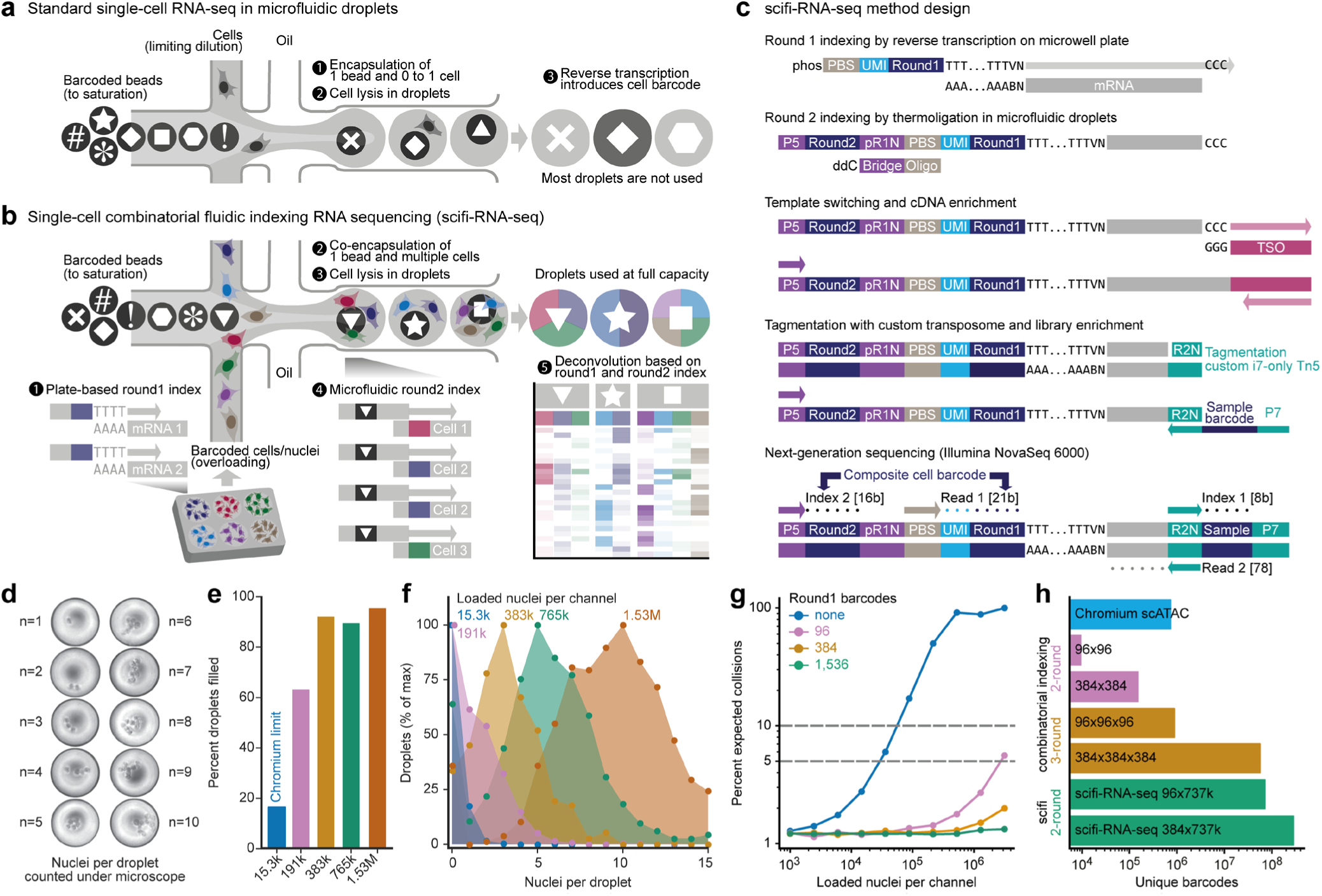
scifi-RNA-seq combines pre-indexing of whole transcriptomes with droplet-based scRNA-seq. **a)** Standard droplet-based scRNA-seq is highly inefficient in its reagent usage. To avoid cell doublets, the single cell suspension is loaded into the microfluidic device at a very low concentration (limiting dilution), leaving most droplets without a cell. Even though the vast majority of droplets contain both a barcoded microbead and reverse transcription reagents, and are thus fully functional, they are not used to generate transcriptome data. Even in droplets that receive one cell, the reagents could in principle support barcoding of additional cells, which adds to the inefficiency. **b)** scifi-RNA-seq uses pre-indexing and droplet overloading to boost the throughput of droplet-based scRNA-seq. Prior to microfluidic indexing, cells or nuclei are permeabilized and their whole transcriptomes are pre-indexed with a well-specific (round1) barcode by reverse transcription on a multiwell plate. Intact cells or nuclei containing differentially barcoded cDNA are randomly mixed and encapsulated in droplets at high concentration using a standard microfluidic droplet generator, such that most droplets contain multiple cells or nuclei (droplet overloading). Inside the droplets, a droplet-specific microfluidic (round2) barcode is added to the pre-indexed cDNA molecules by thermoligation. The combination of round1 and round2 barcodes uniquely identifies single cells. **c)** Detailed method design for scifi-RNA-seq. **d)** By omitting the lysis reagents, intact nuclei can be imaged inside emulsion droplets, confirming the feasibility of droplet overloading using a microfluidic droplet generator (10x Genomics Chromium). Representative droplets containing between one and ten nuclei are shown. **e)** Droplet overloading boosts the percentage of droplets filled with nuclei from 16.4% (maximum loading concentration according to the standard Chromium protocol) to 95.5% (100-fold overloading using 1.53 million nuclei per channel). **f)** Droplet overloading causes the average number of nuclei per droplet to increase in a controlled fashion while maintaining the desired Poisson-like loading distribution. **g)** Expected collision rate as a function of the cell/nuclei loading concentration per channel for defined sets of round1 barcodes. The cell/nuclei fill rate was modelled as a zero-inflated Poisson distribution. **h)** Due to the high number of microfluidic round2 barcodes, two-round scifi-RNA-seq exceeds the barcoding capability of existing three-round combinatorial indexing protocols, while providing a more efficient and greatly simplified workflow.

In droplet-based scRNA-seq, two cells that are loaded into the same droplet will have their transcriptomes labelled with the same cell barcode. As the result, transcripts from these two cells are indistinguishable during data analysis, thus creating a cell doublet that may bias the results (e.g., by creating spurious intermediate cell populations). To minimize the number of cell doublets, the single-cell suspension is loaded into the microfluidic device at a very low concentration, making it unlikely that two cells enter the same droplet. This workaround is sufficient for many applications, but it leads to a major conceptual inefficiency: Most emulsion droplets are fully functional (they contain a barcoded microbead and reverse transcription reagents) but do not receive a cell and therefore do not result in a transcriptome profile (**Fig. 1a**). By design, droplet-based scRNA-seq uses its reagents very inefficiently, which contributes to the method’s high reagent costs and renders it cost-prohibitive for very large studies.

To overcome this conceptual limitation of droplet-based scRNA-seq and to unlock its full potential for massive-scale profiling, we drew inspiration from recently published combinatorial indexing protocols^4-6^. We hypothesized that one round of whole-transcriptome pre-indexing inside intact cells or nuclei, followed by droplet-based scRNA-seq with droplet overloading, could dramatically increase the yield of existing microfluidic droplet generators.

In scifi-RNA-seq (**Fig. 1b-c**), cells or nuclei are permeabilized and their transcriptomes are pre-indexed by reverse transcription in split pools (i.e., in many physically separated bulk aliquots on microwell plates, for example containing 384 pre-indexing (round1) barcodes). Next, cells or nuclei containing pre-indexed cDNA are pooled, randomly mixed, and encapsulated using a standard microfluidic droplet generator, such that most droplets are filled and multiple cells or nuclei occupy the same droplet. Inside the droplets, transcripts are labeled with a microfluidic (round2) barcode. Importantly, neither of the two barcodes is exclusive to a cell, but shared between all cells in the respective reaction compartment (plate well in round1, droplet in round2). Still, because cells or nuclei are randomly mixed between barcoding rounds, the combination of the two barcodes uniquely identifies single cells.

The scifi method is conceptually distinct from cell hashing using DNA-labelled antibodies^7^, lipids^8^, or expressed genetic barcodes^9^. With cell hashing, each cell’s sample origin is encoded by a single index sequence, which provides an effective way of identifying and excluding doublets during data analysis. However, transcripts other than the index sequence cannot be assigned uniquely among multiple cells in the same droplet. In contrast, scifi-RNA-seq can resolve transcripts from overloaded droplets into the respective single-cell transcriptomes thanks to whole-transcriptome pre-indexing. Moreover, our method has relevant practical advantages over multi-round combinatorial indexing protocols, which are very labor-intensive, and result in low transcriptome complexity per cell.

To distinguish our approach from conventional single-cell combinatorial indexing (“sci”) assays using microwell plates, while paying tribute to the elegant design of these methods^4-6^, we refer to the barcoding strategy as “single-cell combinatorial fluidic indexing” (scifi). Here we demonstrate an optimized and validated protocol for scifi-RNA-seq, which makes it possible to obtain up to 150,000 single-cell transcriptomes per channel on the popular Chromium system (10x Genomics)^3^ and more than one million single-cell transcriptomes per microfluidics chip with its eight channels. We achieved high transcriptome complexity per cell, and we successfully validated our method on various cell lines as well as on human primary material, studying T cell activation *ex vivo*.

## Results

### Microfluidic droplet generators can be overloaded with cells or nuclei to enable massive-scale scRNA-seq

Our method is based on the hypothesis that existing microfluidic droplet generators can tolerate substantial overloading (i.e., encapsulation of multiple single cells or nuclei in the same droplet) while maintaining a stable, monodispersed droplet emulsion that does not clog the device. We successfully confirmed this hypothesis for the Chromium system commercialized by 10x Genomics, on which we focused our technology development due to its large user base and the ensuing practical impact. Beyond the Chromium system, our method will also be straightforward to implement on other microfluidic droplet generators including those used in Drop-seq^1^ and inDrop^2^, and it can be adapted to sub-nanoliter well plate assays such as Cyto-seq^10^, Seq-Well^11^, and Microwell-seq^12^.

To test the capacity for droplet overloading on the Chromium system, we prepared a single-nuclei suspension from human Jurkat cells and loaded it at different concentrations. We did not include lysis reagents in these experiments, hence the nuclei inside the droplets stayed intact and could be counted under a light microscope (**Fig. 1d** and **Supplementary Fig. 1a**). We first assessed the maximum recommended loading concentration for the Chromium system, which is 15,300 nuclei per microfluidic channel. Based on 609 representative droplet images, we found that only 16.4% of droplets contained one or more nuclei (mean number of nuclei per droplet: 0.2). Remarkably, even 100-fold overloading (1.53 million nuclei per channel) resulted in a stable droplet emulsion and did not clog the microfluidic system (**Supplementary Fig. 1b**). As we increased the loading concentration, droplet fill rate and number of nuclei per droplet increased in a predictable manner (**Fig. 1e-f**), peaking at a droplet fill rate of 95.5% with an average of 9.6 nuclei per droplet when we loaded 1.53 million nuclei per channel.

Having established that droplet overloading is technically feasible, we sought to quantify the scale of pre-indexing that is required to uniquely resolve single-cell transcriptomes when processing 100,000s to millions of nuclei in a single scifi-RNA-seq experiment. We modeled the number of nuclei per droplet with a zero-inflated Poisson distribution (**Supplementary Fig. 1c-e**), in order to estimate droplet fill rates and the expected percentage of cell doublets for defined numbers of pre-indexing (round1) barcodes as a function of the droplet fill rate (**Fig. 1g**). This analysis indicated that scifi-RNA-seq can uniquely identify large numbers of single cells with relatively few round1 indices, which provides important practical advantages including easier handling, higher sample throughput, and lower setup cost. We independently confirmed this finding by Monte Carlo simulations (**Supplementary Fig. 1f**). Specifically, due to the high complexity of the microfluidic (round2) barcode (for example, the Chromium ATAC-seq reagents that we use in scifi-RNA-seq provide 737,000 distinct microfluidic indices), scifi-RNA-seq with 384-well pre-indexing vastly exceeds the barcoding capacity of three-round (384×384×384) combinatorial indexing (**Fig. 1h**), with the prospect of higher transcriptome complexity per cell and a much easier workflow.

### scifi-RNA-seq boosts the throughput of droplet-based scRNA-seq

The experimental design of scifi-RNA-seq is outlined in **Fig. 1c**, and further technical information including detailed oligonucleotide sequences is provided in **Supplementary Fig. 2** as well as **Supplementary Table 1**. Moreover, a user-friendly and easy to follow step-by-step experimental protocol for scifi-RNA-seq is currently in preparation and will be available upon journal publication.

First, permeabilized cells or nuclei are pre-indexed with barcoded oligo-dT primers by reverse transcription on a multiwell plate (the number of pre-indexing barcodes can be adapted to the required scale; here we used a single 384-well plate). This step labels transcripts with a pre-indexing (round1) barcode specific to each well, while also introducing a unique molecular identifier (UMI) and a primer binding site (PBS) for next-generation sequencing. Upon reaching the end of the transcript, the reverse transcriptase adds untemplated C bases, resulting in a defined end to the cDNA molecules. Second, the cells or nuclei (which still encapsulate the pre-indexed cDNA at this stage) are pooled, washed, filtered, counted, and loaded into the Chromium system at a rate of multiple cells per droplet, along with Chromium gel beads. Inside the droplets, cells or nuclei are lysed and oligonucleotides carrying the microfluidic (round2) barcode are ligated to the cDNA via the 5’-phosphate group of the reverse transcription primer, directed by a complementary 3’-blocked bridge oligo. Efficient ligation is achieved through repeated cycling between denaturation and ligation temperatures with a thermostable ligase. Third, the droplet emulsion is broken, and the 3’-ends of the cDNA molecules are extended by template switching in bulk, followed by cDNA amplification using custom primers. Fourth, double-stranded cDNA is tagmented with a custom i7-only transposome and enriched by PCR, which results in a ready-to-sequence scifi-RNA-seq library.

Additional library barcodes can be introduced in the final step of library preparation, allowing for pooled sequencing of multiple scifi-RNA-seq libraries. Typical fragment distributions are shown in **Supplementary Fig. 3a-b**, and performance metrics for sequencing on the NovaSeq 6000 platform are summarized in **Supplementary Fig. 3c-e**. Notably, by performing only the first step (reverse transcription) on intact nuclei, we achieved excellent nuclei recovery rates for the pre-indexing step (53.3% for cell lines, 41.1% for human primary material, as summarized in **Supplementary Fig. 3f**). Representative images of nuclei containing pre-indexed cDNA are shown in **Supplementary Fig. 3g**).

In addition to the feasibility of droplet overloading (which we confirmed above), a second major feasibility concern was whether the pre-indexed cells and nuclei would withstand the pressures inside the microfluidic droplet generator, given their prior exposure to high-temperature incubations and reagents used in the reverse transcription step. We tested our scifi-RNA-seq protocol with whole cells permeabilized by methanol, with freshly isolated nuclei, and with formaldehyde-fixed frozen nuclei. Results were clearly positive for all three types of input material (**Supplementary Fig. 4**), showing that pre-indexed cells and nuclei are indeed compatible with subsequent fluidic indexing. Although cells and nuclei performed similarly in this comparison, we decided to focus our further experiments on nuclei, as their small size makes it possible to achieve very high droplet fill rates and their homogenous RNA content may result in more balanced sequencing coverage for massive-scale profiling.

To evaluate the performance of scifi-RNA-seq as a function of droplet overloading, we loaded 15,300, 383,000, or 765,000 pre-indexed nuclei into a single channel of the Chromium system, and we determined the inflection point in the distribution of UMIs across single-cell barcodes (**Fig. 2a**), which separates nuclei from noise in droplet-based scRNA-experiments. We found that the number of recovered single-cell transcriptomes scaled linearly with the number of loaded nuclei. Furthermore, the barcoding allowed us to determine the exact number of individual nuclei (as identified by round1 barcodes) in each droplet (as identified by round2 barcodes) based on the sequencing data (**Fig. 2b**). The average number of nuclei inside each droplet increased in a controlled fashion, peaking at an average of 4.4 nuclei per droplet when 765,000 nuclei were loaded. Taken together, our results indicate that scifi-RNA-seq recovers nuclei with comparable efficiency over a wide range of loading concentrations, confirming our model based on the optical inspection of the droplet emulsion (**Supplementary Fig. 1c-f**).

**Figure 2:**
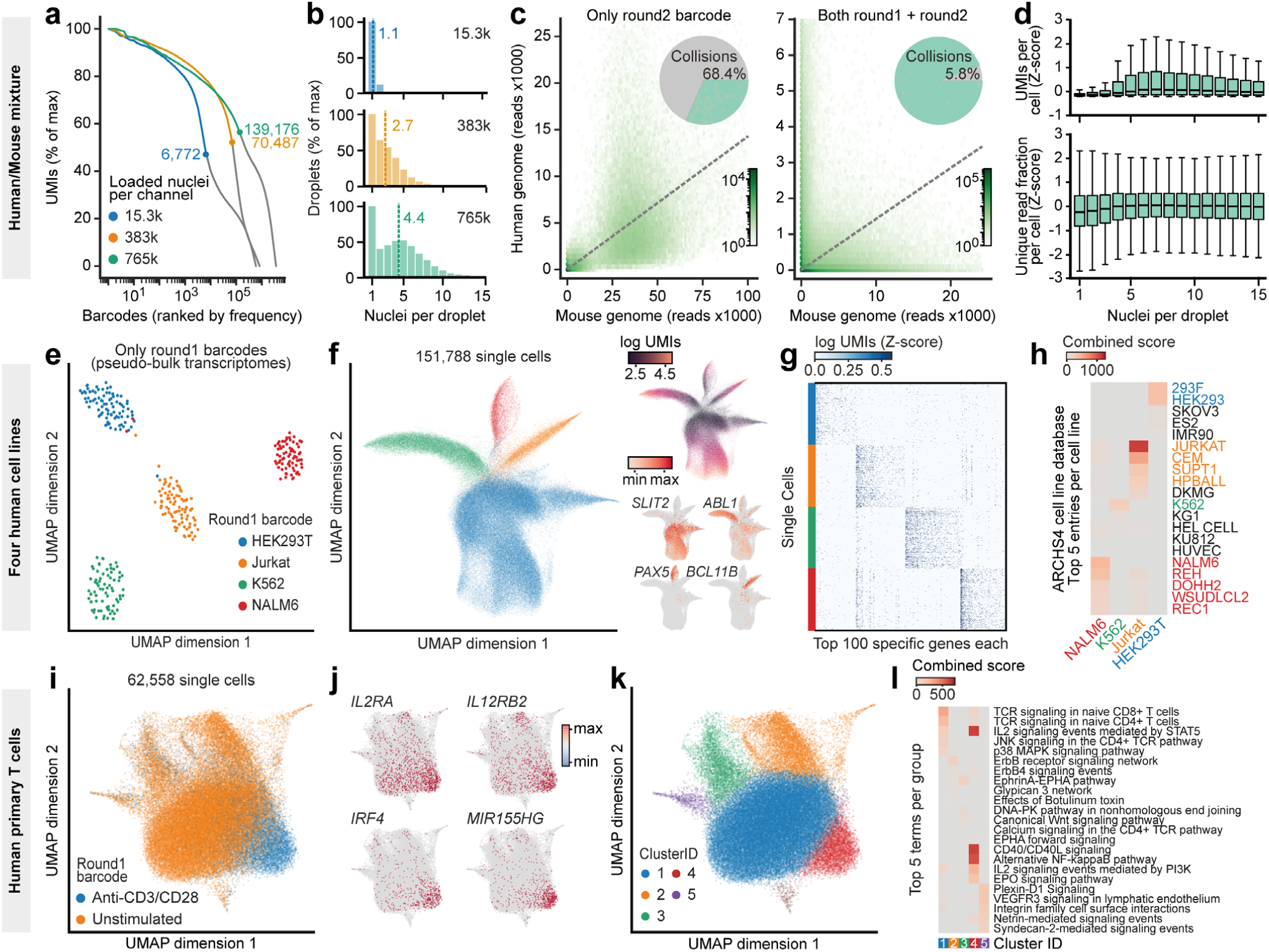
scifi-RNA-seq produces high-quality single-cell transcriptomes for cell lines and primary samples. **a)** A mixture of human and mouse nuclei (Jurkat and 3T3, respectively) was processed with scifi-RNA-seq, loading 15,300, 383,000, and 765,000 nuclei into single microfluidic channels of the Chromium device. Plotting all detected barcodes ranked by frequency against the number of unique molecular identifiers (UMIs) per barcode identifies a characteristic inflection point that separates nuclei (left, colored lines) from background noise (right, grey lines). **b)** Distribution of the number of nuclei (round1 indices) per droplet (round2 barcode) for increasing nuclei loading concentrations. The average number of nuclei per droplet and nuclei loading concentration per channel are indicated. **c)** The round1 transcriptome index can deconvolute multiple nuclei per droplet into the respective single-cell transcriptomes. 765,000 pre-indexed nuclei from a mixture of human (Jurkat) and mouse (3T3) cells were processed in a single microfluidic channel and demultiplexed based on the microfluidic round2 barcode only (left plot), or based on the combination of round1 and round2 barcodes (right plot). The percentages of detected inter-species collisions are shown by the pie charts. The dashed line indicates the expected 1:1 mixture ratio, accounting for the relative transcript content of cells for each species. **d)** UMIs per cell and fraction of unique reads per cell are plotted against the number of nuclei contained in the respective droplet, showing no deterioration in the single-cell transcriptome complexity when many cells co-occupy the same droplet. This analysis is based on the largest human/mouse mixing experiment with 765,000 nuclei per microfluidic channel. **e)** Four human cell lines (HEK293T, Jurkat, K562, NALM6) were processed with scifi-RNA-seq, using defined sets of round1 barcodes for each cell line. Considering only round1 barcodes, the dataset gives rise to averaged pseudo-bulk RNA-seq profiles of the cell lines, which are plotted here. **f)** 151,788 single-cell transcriptomes derived from the human cell line mixture are displayed in a 2D projection using the UMAP algorithm and colored by round1 barcodes corresponding to cell lines (left), UMIs per cell (top right), or marker gene expression (bottom right). **g)** Heatmap showing single-cell expression levels for the top 100 most specific genes for each cell line. We randomly sampled an equal number of single-cell transcriptomes per cell line without filtering for transcriptome quality. **h)** Gene set enrichment analysis of differentially expressed genes clearly identifies the cell lines. Colors indicate the assayed cell lines as well as closely related cell lines in the ARCHS4 database. **i)** Human primary T cells with or without T cell receptor stimulation were processed using scifi-RNA-seq, and the single-cell transcriptomes are displayed in a UMAP projection (color-coded by stimulation state). **j)** Expression levels of four genes induced by TCR stimulation overlaid on the UMAP projection. **k)** UMAP projection with single cells colored by clusters assigned by graph-based clustering using the Leiden algorithm. **l)** Gene set enrichment analysis for the differentially expressed genes in each cluster according to panel k.

This test dataset, which was based on a 1:1 mixture of human and mouse cell lines (Jurkat and 3T3, respectively), also allowed us to validate our pre-indexing strategy for the correct assignment of transcripts to single cells. To that end, we compared the number of human-mouse cell doublets based on the microfluidic (round2) barcode only with the number of such doublets based on the combination of pre-indexing (round1) and microfluidic (round2) barcodes. As expected for a loading rate of 765,000 nuclei per channel, almost all droplets contained both human and mouse cells (**Fig. 2c**, left panel), but we could resolve the vast majority of these doublets when considering both the round1 and round2 barcode (**Fig. 2c**, right panel). As expected, the striking effect of pre-indexing was seen only when the droplet generator was overloaded, while the microfluidic round2 barcode alone was sufficient for minimizing cell doublets at a standard loading rate of 15,300 nuclei per channel (**Supplementary Fig. 5a**).

Finally, the dataset allowed us to conclusively resolve a third feasibility concern for scifi-RNA-seq – whether the reagents in each droplet would be sufficient for effective barcoding of the transcriptomes from multiple nuclei. When plotting UMI counts and fractions of unique reads per cell against the number of nuclei per droplet (**Fig. 2d**), we observed no trend toward lower transcriptome complexity in droplets containing up to 15 individual nuclei, strongly suggesting that reagents for droplet-based indexing are not a limiting factor in scifi-RNA-seq.

### scifi-RNA-seq supports massive-scale scRNA-seq on cell lines and primary samples

Having established the general feasibility of scifi-RNA-seq, we sought to benchmark the method’s performance for different cell types. To this end, we first prepared nuclei from four human cell lines with unique characteristics (HEK293T, Jurkat, K562, NALM-6) and subjected these cells to scifi-RNA-seq with 384-well pre-indexing. Each cell line was assigned a specific set of pre-indexing (round1) barcodes, allowing us to demonstrate the method’s inherent support for multiplexing thousands of distinct samples (when using more than one 384-well plate) in a single experiment (**Fig. 2e**). After pre-indexing, samples were pooled and 383,000 nuclei were loaded into a single microfluidic channel of the Chromium system. This experiment resulted in 151,788 single-cell transcriptomes passing quality control, which constitutes a 15-fold increase over the output of the standard Chromium protocol. These single-cell transcriptomes clustered into four different point clouds corresponding to the four cell lines (**Fig. 2f**, left), with clear separation between cell lines for all but the most shallowly sequenced cells (**Fig. 2f**, top right). The transcriptome data quality was high enough that even single marker genes such as *ABL1* for K562 and *PAX5* for NALM-6 were able to distinguish between the different cell lines (**Fig. 2f**, bottom right), and the clustering was immediately apparent from the expression of the top-100 differential genes per cell line without filtering for high-coverage transcriptomes (**Fig. 2g**). Enrichment analysis against publicly available gene expression signatures accurately identified the corresponding cell lines based on the single-cell transcriptome data (**Fig. 2h**). In summary, profiling of four human cell lines with ∼150,000 single-cell transcriptomes showed that scifi-RNA-seq yields single-cell transcriptomes of high quality for cells cultured *in vitro*, with potential future applications in drug screens.

Next, we assessed how well scifi-RNA-seq performs on human primary cells, in a model where the differences in gene expression are more subtle. We purified human CD3+ T cells from the peripheral blood of three healthy donors and maintained the isolated T cells in short-term culture using human T cell medium containing IL-2. Half of the cells were T cell receptor (TCR) stimulated with anti-CD3/CD28 activator beads, while the other half were left untreated. Nuclei were fixed with formaldehyde and cryopreserved for easier handling. All samples were processed in a single scifi-RNA-seq experiment, using the pre-indexing step to uniquely barcode both the donors and the stimulation states. We obtained 62,558 single-cell transcriptomes that passed quality control.

Unsupervised analysis of the single-cell transcriptomes was dominated by TCR stimulation state (**Fig. 2i**). Our data identified characteristic markers of T cell activation (**Fig. 2j**) including *IL2RA* (CD25) and *IL12RB2* (which encode cytokine receptors) as well as *IRF4* and *MIR155HG* (which are known regulators of T cell biology). Graph-based clustering identified five groups of cells characterized by similar transcriptome profiles (**Fig. 2k**), which we annotated by gene set analysis (**Fig. 2l**). We found significant and cluster-specific enrichment of signaling path-ways relevant to T cell activation, including signaling via the TCR, IL-2, CD40L, and NFκB. In summary, these results show that scifi-RNA-seq is applicable to primary cells and able to detect modest transcriptional differences between cell states, with potential future applications in immuno-oncology and cell atlas projects.

## Discussion

Here we described the “single-cell combinatorial fluidic indexing” (scifi) concept and its initial implementation in the scifi-RNA-seq assay, providing a method for massive-scale scRNA sequencing that is efficient, flexible, and easy-to-use. scifi-RNA-seq is straightforward to set up, only requiring standard laboratory equipment and access to a microfluidic droplet generator such as the 10x Genomics Chromium system (optionally via shipment of fixed nuclei to a core facility). A user-friendly and easy to follow step-by-step experimental protocol for scifi-RNA-seq is currently in preparation and will be available upon journal publication.

Our method combines the scalability of combinatorial indexing with the efficiency and ease-of-use of droplet generators in a straightforward two-stage workflow. Pre-indexing on 384-well plates (round1) boosts the throughput of subsequent droplet-based scRNA-seq by enabling droplet overloading. We demonstrated a 15-fold increase in throughput, with scope for even larger scale given that we were able to load 1.53 million nuclei into a single channel of the Chromium system. The microfluidic (round2) indexing with its high number of barcodes distinguishes our method from existing combinatorial indexing protocols, where at least three (and sometimes up to five) rounds of plate-based indexing are needed to achieve high throughput. With just two rounds of indexing and only the reverse transcription done inside intact cells or nuclei, scifi-RNA-seq provides an easier, faster, and more efficient workflow for massive-scale transcriptome profiling, making it a method of choice for many biological applications.

We expect scifi-RNA-seq to be most immediately useful for two application areas: Cell atlas projects pursuing the single-cell characterization of complex tissues, organs, or entire organisms^13, 14^; and massive-scale perturbation studies including drug screens^15^ and single-cell CRISPR screens (e.g., using CROP-seq^16^, Perturb-seq^17, 18^, or CRISP-seq^19^). Both applications profit not only from the massive scale that scifi-RNA-seq provides, but also from its cost-efficiency and built-in support for multiplexing. Compared to droplet-based scRNA-seq, where library preparation costs may account for 80% of the total spending on consumables, the actual sequencing costs become the only limiting factor for very large scifi-RNA-seq experiments. The massive scale of scifi-RNA-seq benefits applications that require very deep profiling of complex samples (e.g., to discover rare stem cell populations in human tissue, cancer cells in peripheral blood, or intermediate cell states in cell reprogramming experiments).

Moreover, scifi-RNA-seq facilitates projects with large numbers of individual samples or conditions. Because the pre-indexing is performed on 96-well or 384-well plates, where each well receives a unique barcode, it is straight-forward to multiplex hundreds or even thousands of individual samples in a single scifi-RNA-seq experiment – for example measuring the transcriptome response to a large number of genetic or pharmacological perturbations in cell lines, or processing tissue samples from large epidemiological cohorts. Combined with efficient PCR-based guide RNA enrichment, scifi-RNA-seq could also become the method of choice for single-cell CRISPR screens, where very high cell numbers are needed for large-scale genetic screens with transcriptome readout. Finally, scifi-RNA-seq may even provide a cost-effective method for bulk transcriptome profiling in very large sample collections, exploiting the high multiplexing capability of pre-indexing combined with the low reagent consumption in single cells and subsequent derivation of pseudo-bulk transcriptome profiles from single-cell data.

While our initial demonstration of scifi-RNA-seq focuses on the widely available 10x Genomics Chromium system, scifi-RNA-seq will be straightforward to adapt to other microfluidic droplet generators, and it can be combined with alternative methods for large-scale scRNA-seq such as sub-nanoliter well plates. More generally, the scifi concept is not limited to scRNA-seq, but could also be used to boost the throughput of other single-cell assays such as single-cell ATAC-seq^20, 21^, whole genome sequencing^22^, DNA-methylation analysis^23^, and chromatin conformation capture by Hi-C^24^. With our thermoligation barcoding reaction optimized for use inside emulsion droplets, we provide a versatile enzymatic approach to support these and other applications, which may in the future include multi-omics protocols^25^ such as combined RNA-seq and ATAC-seq in single cells^26-28^.

In conclusion, we expect the scifi concept and its implementation in the scifi-RNA-seq protocol to provide an easy-to-use, cost-effective, and broadly useful method for those applications in single-cell biology that profit from massive throughput and from the ability to multiplex many samples.

## Online Methods

### Measurement of bead fill rates

We loaded the Chromium Single Cell E Chip (10x Genomics #2000121) with 80 µl of 1x Nuclei Buffer (10x Genomics #2000153) into inlet 1, 40 µl of Single Cell ATAC Gel Beads (10x Genomics #2000132) into inlet 2, and 240 µl of Partitioning Oil (10x Genomics #220088) into inlet 3. Because we omitted Reducing Agent B, the gel beads remained intact throughout the microfluidic run, such that they could be visualized inside the emulsion droplets using a standard light microscope. Our fill rate calculations are based on a total of 1,265 droplets that were manually counted from microscopic images.

### Testing of the nuclei loading capacity

Human Jurkat cells (clone E6-1) were cultured in RPMI medium (Gibco #21875-034) supplemented with 10% FCS (Sigma) and penicillin-streptomycin (Gibco #15140122). Fresh nuclei were isolated as described below. We prepared samples of 15,300, 191,000, 383,000, 765,000, and 1,530,000 nuclei, added 1.5 µl of Reducing Agent B (10x Genomics #2000087) and 1x Nuclei Buffer (10x Genomics #2000153) to a total volume of 80 µl. This buffer did not contain detergents, hence the nuclei remained intact during the microfluidic run and could be visualized inside the emulsion droplets with a standard light microscope. At the same time, Reducing Agent B dissolves the gel beads, which might otherwise obstruct the view. The Single Cell E Chip (10x Genomics #2000121) was loaded as follows: 75 µl of nuclei suspension at the indicated loading concentrations into inlet 1, 40 µl of Single Cell ATAC Gel Beads (10x Genomics #2000132) into inlet 2, and 240 µl of Partitioning Oil (10x Genomics #220088) into inlet 3. To image the resulting droplets, 15 µl of Partitioning Oil were pipetted onto a glass slide, followed by 5 µl of emulsion droplets, and images were taken at 10x magnification. Nuclei were manually counted from microscopic images for an average of 653 droplets per loading concentration.

### Preparation of permeabilized cell suspension

A total of 5 million cells were washed with 10 ml of ice-cold 1x PBS (Gibco #14190-094) with centrifugation (300 rcf, 5 min, 4 °C) and fixed in 5 ml of ice-cold methanol (Fisher Scientific #M/4000/17) at -20 °C for 10 min. After two additional washes (centrifugation: 300 rcf, 5 min, 4 °C) with 5 ml of ice-cold PBS-BSA-SUPERase (1x PBS supplemented with 1% w/v BSA (Sigma #A8806-5) and 1% v/v SUPERase-In RNase Inhibitor (Thermo Fisher Scientific #AM2696)) permeabilized cells were resuspended in 200 µl of ice-cold PBS-BSA-SUPERase, and filtered through a cell strainer (40 µM or 70 µM depending on the cell size). We then used 5 µl of the sample for cell counting in duplicates on a CASY device (Schärfe System) and diluted to 5,000 cells per µl with ice-cold PBS-BSA-SUPERase. We immediately proceeded with the reverse transcription step.

### Preparation of fresh nuclei suspension

A total of 5 million cells were washed with 10 ml of ice-cold 1x PBS (Gibco #14190-094) with centrifugation (300 rcf, 5 min, 4 °C). Nuclei were prepared by resuspending cells in 500 µl of ice-cold Nuclei Preparation Buffer (10 mM Tris-HCl pH 7.5 (Sigma #T2944-100ML), 10 mM NaCl (Sigma #S5150-1L), 3 mM MgCl2 (Ambion #AM9530G), 1% w/v BSA (Sigma #A8806-5), 1% v/v SUPERase-In RNase Inhibitor (20 U/µl, Thermo Fisher Scientific #AM2696), 0.1% v/v Tween-20 (Sigma #P7949-500ML), 0.1% v/v IGEPAL CA-630 (Sigma #I8896-50ML), 0.01% v/v Digitonin (Promega #G944A)), followed by 5 min of incubation on ice. Lysis of the plasma membrane was stopped by adding 5 ml of ice-cold Nuclei Wash Buffer (10 mM Tris-HCl pH 7.5, 10 mM NaCl, 3 mM MgCl2, 1% w/v BSA, 1% v/v SUPERase-In RNase Inhibitor, 0.1% v/v Tween-20). Nuclei were collected by centrifugation (500 rcf, 5 min, 4 °C), resuspended in 200 µl of ice-cold PBS-BSA-SUPERase (1x PBS supplemented with 1% w/v BSA and 1% v/v SUPERase-In RNase Inhibitor) and filtered through a cell strainer (40 µM or 70 µM depending on the cell size). We then used 5 µl of the sample for cell counting in duplicates on a CASY device (Schärfe System) and diluted to 5,000 nuclei per µl with ice-cold PBS-BSA-SUPERase. We immediately proceeded with the reverse transcription step.

### Preparation of fixed nuclei suspension

A total of 5 million primary cells were washed with 10 ml of ice-cold 1x PBS (Gibco #14190-094) with centrifugation (300 rcf, 5 min, 4 °C). Nuclei were prepared by resuspending cells in 500 µl of ice-cold Nuclei Preparation Buffer without Digitonin and without Tween-20 (10 mM Tris-HCl pH 7.5 (Sigma #T2944-100ML), 10 mM NaCl (Sigma #S5150-1L), 3 mM MgCl2 (Ambion #AM9530G), 1% w/v BSA (Sigma #A8806-5), 1% v/v SUPERase-In RNase Inhibitor (Thermo Fisher Scientific #AM2696), 0.1% v/v IGEPAL CA-630 (Sigma #I8896-50ML)), followed by 5 min of incubation on ice. Lysis of the plasma membrane was stopped by addition of 5 ml of Nuclei Wash Buffer without Tween-20 (10 mM Tris-HCl pH 7.5, 10 mM NaCl, 3 mM MgCl2, 1% w/v BSA, 1% v/v SUPERase-In RNase Inhibitor). Nuclei were collected by centrifugation (500 rcf, 5 min, 4 °C), and fixed in 5 ml of ice-cold 1x PBS containing 4% Formaldehyde (Thermo Fisher Scientific #28908) for 15 min on ice. Fixed nuclei were collected (500 rcf, 5 min, 4 °C), the pellet was resuspended in 1.5 ml of ice-cold Nuclei Wash Buffer without Tween-20 and transferred to a 1.5 ml tube. After one more wash with 1.5 ml of ice-cold Nuclei Wash Buffer without Tween-20 (500 rcf, 5 min, 4 °C), fixed nuclei were resuspended in 200 µl of Nuclei Wash Buffer without Tween-20, snap-frozen in liquid nitrogen and stored at -80 °C.

For scifi-RNA-seq, frozen samples were thawed in a 37 °C water bath for exactly 1 min, and immediately placed on ice. Following centrifugation (500 rcf, 5 min, 4 °C), fixed nuclei were resuspended in 250 µl of ice-cold Permeabilization Buffer (10 mM Tris-HCl, 10 mM NaCl, 3 mM MgCl2, 1% w/v BSA, 1% v/v SUPERase-In RNase Inhibitor, 0.01% v/v Digitonin (Promega #G944A), 0.1% v/v Tween-20 (Sigma cat. no P7949-500ML)). After 5 min of incubation on ice, 250 µl of Nuclei Wash Buffer without Tween-20 were added per sample, and nuclei were collected (500 rcf, 5 min, 4 °C). After one more wash with 250 µl of Nuclei Wash Buffer without Tween-20, nuclei were taken up in 100 µl of 1x PBS containing 1% w/v BSA and 1% v/v SUPERase-In RNase Inhibitor. We used 5 µl for cell counting in duplicates on a CASY device (Schärfe Systems) and diluted to 5,000 nuclei per µl with PBS-BSA-SUPERase. We immediately proceeded with the reverse transcription step.

### Isolation of primary human T cells

Peripheral blood from healthy donors was obtained from the Austrian Red Cross as blood packs with buffered sodium citrate as anti-coagulant. The study complied with all relevant ethical regulations for working with human primary samples. Informed consent was obtained from all sample donors. The study was approved by the ethical committees of the contributing institutions (Austrian Red Cross, Medical University of Vienna). For each donor, we prepared T cells from 3x 15 ml of peripheral blood, according to the following protocol. 15 ml of peripheral blood were mixed with 750 µl of RosetteSep Human T Cell Enrichment Cocktail (Stemcell #15061). After 10 min of incubation at room temperature, the sample was diluted by addition of 15 ml 1x PBS (Gibco #14190-094) containing 2% v/v FCS (Sigma). SepMate tubes (Stemcell #86450) were loaded with 15 ml of Lymphoprep density gradient medium (Stemcell #07851) and the blood sample was poured on top. After centrifugation (1,200 rcf, 10 min, room temperature, brake set to 9), the supernatant was transferred to a fresh 50 ml tube, topped up to 50 ml with 1x PBS containing 2% FCS, and centrifuged (1200 rcf, 10 min, room temperature, brake set to 3). After one additional wash with 50 ml of 1x PBS containing 2% FCS (1200 rcf, 10 min, room temperature, brake set to 3), T cells were resuspended in 10 ml of 1x PBS containing 2% FCS, filtered through a 40 µM cell strainer, and counted using a CASY device (Schärfe Systems). For accurate cell counting, it was important to exclude contaminating erythrocytes, which will be lysed during the subsequent nuclei preparation.

### Anti-CD3/CD28 stimulation of human T cells

Freshly isolated primary human T cells were resuspended at a density of 1 million cells per ml in Human T Cell Medium (OpTmizer medium (Thermo Fisher #A1048501) containing 1/38.5 volumes of OpTmizer supplement, 1x GlutaMax (Thermo Fisher #35050061), 1x Penicillin/Streptomycin (Thermo Fisher #15140122), 2% heat-inactivated human AB serum (Fisher Scientific #MT35060CI), 10 ng/ml of recombinant human IL-2 (PeproTech #200-02)). The culture was split into two flasks, and one was treated with Human T-Activator CD3/CD28 Dynabeads (25 µl beads per 1 million cells, Thermo Fisher #11131D). After 16 hours, we prepared formaldehyde-fixed nuclei and snap-froze the nuclei suspension as described above.

### Assembly and validation of custom i7-only transposome for scifi-RNA-seq

Oligonucleotides Tn5-top_ME and Tn5-bottom_Read2N were synthesized by Sigma Aldrich (sequences are provided in **Supplementary Table 1**) and reconstituted in EB buffer (Qiagen #19086) at 100 µM. We mixed 22.5 µl of each oligonucleotide with 5 µl of 10x Oligonucleotide Annealing Buffer (10 mM Tris-HCl (Sigma #T2944-100ML), 500 mM NaCl (Sigma #S5150-1L), 10 mM EDTA (Invitrogen #AM9260G)) and annealed them in a thermocycler (95 °C for 3 min, 70 °C for 3 min, ramp to 25 °C at -2 °C per minute). The annealing reaction was then diluted by addition of 180 µl of water. At this point, the diluted oligonucleotide cassette can be aliquoted and frozen for future transposome assemblies. To load the Tn5 transposase, we mixed 20 µl of diluted oligonucleotide cassette from the previous step with 20 µl of 100% glycerol (Sigma #G5516-100ML) and 10 µl of EZ-Tn5 Transposase (Lucigen #TNP92110), and incubated for 30 min at 25 °C in a thermocycler. The resulting 50 µl of assembled transposome can be stored at -20 °C for at least one month.

Tagmented DNA flanked by two Illumina i7 adapters is suppressed in standard PCR reactions due to competition between intramolecular annealing and primer binding, as described previously^29^. However, the enzymatic activity of the custom i7-only transposome can still be tested in a negative qPCR assay. Briefly, a defined PCR product was subjected to one tagmentation reaction and one no-enzyme control reaction, and both samples were re-amplified with the same primers by qPCR. Since the tagmentation fragments the PCR product, the corresponding reaction should yield higher Ct values. The tagmentation efficiency can then be calculated from the shift of Ct values:

Tagmentation efficiency [%] = 100 − (100 / (2 ^ (average Ct tagmentation − average Ct no-enzyme control)))

The PCR product for testing the enzymatic activity was produced as follows. Oligonucleotides pUC19-FWD and pUC19-REV were synthesized by Sigma Aldrich (sequences are provided in **Supplementary Table 1**) and reconstituted in EB buffer (Qiagen #19086) at 100 µM. Next, a 1,961 bp PCR product was generated by mixing 128.7 µl of water, 33 µl of 50 pg/µl pUC19 plasmid (NEB #N3041S), 1.65 µl each of primers pUC19-FWD and pUC19-REV (100 µM) combined with 165 µl of 2x Q5 HotStart High-Fidelity Master Mix (NEB #M0494L). The resulting 6.6x master mix was distributed into a tube strip (six reactions of 50 µl) and amplified in a thermocycler: 98 °C for 30 s; 31x (98 °C for 10 s, 68 °C for 30 s, 72 °C for 1 min), 72 °C for 2 min, storage at 12 °C. To each 50 µl PCR reaction, we added 6.25 µl of 10x CutSmart Buffer and 6.25 µl of DpnI (NEB #R0176L) and incubated at 37 °C for 1 hour to digest the PCR template plasmid. The six PCR reactions were pooled and cleaned with the QiaQuick PCR Purification Kit (Qiagen #28106) using two columns and eluting with 30 µl of EB buffer per column. Eluates were pooled, and the purity of the PCR fragment was checked on a 1% agarose gel containing ethidium bromide. We then measured the concentration of dsDNA with a Qubit HS assay (Thermo Fisher Scientific #Q32854), and diluted the PCR product to 25 ng/µl with EB buffer.

Based on the resulting PCR product, tagmentation reactions were set up by mixing 2 µl of 25 ng/µl of the PCR product, 7 µl of ATAC Buffer (10x Genomics #2000122), and either 6 µl of custom i7-only transposome (tagmentation reaction) or 6 µl of water (no-enzyme control reaction). After 60 min of incubation at 37 °C, the Tn5 enzyme was stripped from the DNA by addition of 1.75 µl of 1% SDS solution (Sigma #71736-100ML) followed by incubation at 70 °C for 10 min. The two reactions were diluted 1/100 with EB buffer, and qPCR reactions were set up in triplicates: 2 µl of 1/100-diluted reaction, 10 µl of 2x GoTaq qPCR Master Mix (Promega #A600A), 0.1 µl each of 100 µM pUC19-FWD and pUC19-REV primers and 7.8 µl of water. qPCR reactions were incubated as follows: 95 °C for 2 min, 40x (95 °C for 30 s, 68 °C for 30 s, 72 °C for 2 min, plate reading).

### Detailed scifi-RNA-seq protocol description

#### Reverse Transcription

384 indexed reverse transcription primers were synthesized by Sigma Aldrich and obtained at 100 µM concentration in EB Buffer on 96-well plates (sequences are provided in **Supplementary Table 1**). 384-well plates with barcoded oligo-dT primers were prepared prior to the experiment and stored at -20 °C (1 µl of 25 µM per well). 10,000 permeabilized cells or nuclei (2 µl of a 5,000/µl suspension) were added to the pre-dispensed primers in each well, and the assignment of samples to wells was recorded. The plate was incubated for 5 min at 55 °C to resolve RNA secondary structures, then placed immediately on ice to prevent their re-formation. Per well, a mix of 3 µl nuclease-free water, 2 µl 5x Reverse Transcription Buffer, 0.5 µl of 100 mM DTT (freshly diluted from Sigma #646563-10x.5ML), 0.5 µl of 10 mM dNTPs (Thermo Fisher Scientific #R0193), 0.5 µl of RNaseOUT RNase inhibitor (40 U/ml, Thermo Fisher Scientific #10777019), and 0.5 µl of Maxima H Minus Reverse Transcriptase (200 U/ml, Thermo Fisher Scientific #EP0753) were added. The reverse transcription was incubated as follows (with heated lid set to 60 °C): 50 °C for 10 min, 3 cycles of [8 °C for 12 s, 15 °C for 45 s, 20 °C for 45 s, 30 °C for 30 s, 42 °C for 2 min, 50 °C for 3 min], 50 °C for 5 min, store at 4 °C.

#### Cell/Nuclei recovery and pooling

Processed cells/nuclei were recovered from the 384-well plate and pooled in one 15 ml tube per plate on ice. Wells were washed with ice-cold 1x PBS containing 1% BSA, which was transferred to the same tube for maximum recovery. The volume was topped up to 15 ml with 1x PBS containing 1% BSA, and nuclei were collected (500 rcf, 5 min, 4 °C). The resulting pellet was resuspended in 1.0 ml of 1x Ampligase Reaction Buffer (Lucigen #A0102K), filtered through a cell strainer (40 µm or 70 µm depending on the cell/nuclei size) into a 1.5 ml tube, and centrifuged (500 rcf, 5 min, 4 °C). The supernatant was removed almost completely, and the tube was centrifuged briefly (500 rcf, 30 s, 4 °C) to collect the remaining liquid at the bottom of the tube. Typically, this resulted in ∼10 µl of a highly concentrated suspension, which was diluted 1:200 with 1x Ampligase Buffer and counted in a Fuchs Rosenthal counting chamber (Incyto #DHC-F01). The desired number of cells/nuclei was brought to a volume of 15 µl with 1x Ampligase Reaction Buffer (Lucigen #A0102K).

#### Microfluidic thermoligation barcoding

Unused channels in the Chromium Chip E (10x Genomics #2000121) were filled with 75 µl (inlet 1), 40 µl (inlet 2), and 240 µl (inlet 3) of 50% glycerol solution (Sigma #G5516-100ML). Right before loading the chip, a mix of 47.4 µl nuclease-free water, 11.5 µl of 10x Ampligase Reaction Buffer (Lucigen #A0102K), 2.3 µl of 100 U/µl Ampligase (Lucigen #A0102K), 1.5 µl of Reducing Agent B (10x Genomics #2000087), and 2.3 µl of 100 µM Bridge Oligo (sequence provided in **Supplementary Table 1**) was added per sample. The microfluidic chip was loaded with 75 µl of cells or nuclei in thermoligation mix (inlet 1), 40 µl of Single Cell ATAC Gel Beads (inlet 2, 10x Genomics #2000132), and 240 µl of Partitioning Oil (inlet 3, 10x Genomics #220088) and run on the Chromium system. For thermoligation barcoding, the droplet emulsion was incubated as follows (heated lid set to 105 °C, volume set to 100 µl): 12 cycles of [98 °C for 30 s, 59 °C for 2 min], storage at 15 °C. The emulsion was broken by addition of 125 µl Recovery Agent (10x Genomics #220016), and 125 µl of the pink oil phase were removed by pipetting. The remaining sample was mixed with 200 µl of Dynabead Cleanup Master Mix (per reaction: 182 µl Cleanup Buffer (10x Genomics #2000088), 8 µl Dynabeads MyOne Silane (Thermo Fisher Scientific #37002D), 5 µl Reducing Agent B (10x Genomics #2000087), 5 µl of nuclease-free water). After 10 min of incubation at room temperature, samples were washed twice with 200 µl of freshly prepared 80% ethanol (Merck #603-002-00-5) and eluted in 40.5 µl of EB Buffer (Qiagen #19086) containing 0.1% Tween (Sigma #P7949-500ML) and 1% v/v Reducing Agent B. Bead clumps were sheared with a 10 µl pipette or needle. 40 µl of the sample were transferred to a fresh tube strip and subjected to a 1.0x cleanup with SPRIselect beads (Beckman Coulter #B23318), eluting in 22 µl of EB Buffer.

#### Template switching

20 µl of sample from the previous step were mixed with 10 µl of 5x Reverse Transcription Buffer, 10 µl of Ficoll PM-400 (20%, Sigma #F5415-50ML), 5 µl of 10 mM dNTPs (Thermo Fisher Scientific #R0193), 1.25 µl of Recombinant Ribonuclease Inhibitor (Takara #2313A), 1.25 µl of 100 µM Template Switching Oligo (sequence provided in **Supplementary Table 1**), and 2.5 µl of Maxima H Minus Reverse Transcriptase (200 U/ml, Thermo Fisher Scientific #EP0753). The template switching reaction was incubated for 30 min at 25 °C, 90 min at 42 °C, storage at 4 °C, and cleaned with a 1.0x SPRI cleanup, eluting in 17 µl of EB buffer.

#### cDNA enrichment

15 µl of the above sample were mixed with 33 µl of nuclease-free water, 50 µl of NEBNext High-Fidelity 2x PCR Master Mix (NEB #M0541S), 0.5 µl of 100 µM Partial P5 primer, 0.5 µl of 100 µM TSO Enrichment Primer (sequences provided in **Supplementary Table 1**), and 1 µl of 100x SYBR Green in DMSO (Life Technologies #S7563). cDNA was amplified in a thermocycler as follows: 98 °C for 30 s, cycle until fluorescent signal >1000 RFU [98 °C for 20 s, 65 °C for 30 s, 72 °C for 3 min], 72 °C for 5 min in another thermocycler, storage at 4 °C. cDNA was cleaned by one 0.8x SPRI cleanup followed by a 0.6x SPRI cleanup, quantified with a Qubit HS assay (ThermoFisher Scientific #Q32854), and 1.5 ng were checked on a Bioanalyzer High-Sensitivity DNA chip (Agilent #5067-4626 and 5067-4627).

#### Tagmentation

To achieve maximum library complexity, the entire sample was processed in multiple tagmentation reactions with 1 ng input each. cDNA was diluted to 0.2 ng/µl with nuclease-free water and 5 µl (1 ng) per reaction were distributed into a 96-well plate on ice. A mix of 11.25 µl nuclease-free water, 5 µl of 5x Tn5 Reaction Buffer (50 mM TAPS (Sigma #T9659-100G), 25 mM MgCl_2_ (Ambion #AM9530G), pH adjusted to 8.5, sterile-filtered), 2.5 µl of Dimethylformamide (Sigma #D4551-250ML), and 1.25 µl of freshly diluted i7-only transposome (prepared as described above and diluted 1:4.5 in Tn5 Dilution Buffer (50 mM Tris-HCl pH 7.5 (Sigma #T2944-100ML), 100 mM NaCl (Sigma #S5150-1L), 0.1 mM EDTA (Invitrogen #AM9260G), 50% glycerol (Sigma #G5516-100ML), 0.1% Triton-X100 (Sigma #X100-100ML), 1 mM freshly added DTT (Sigma #646563-10x.5ML)) was added. Reactions were incubated for 10 min at 55 °C, then cooled for 1 min on ice. The enzyme was inactivated by addition of 2.5 µl of 1% SDS (Sigma #71736-100ML) for 5 min at room temperature. Next, the volume was brought to 50 µl and the fragmented cDNA was purified with a 1.0x SPRI cleanup, eluting in 17 µl of EB buffer (Qiagen #19086).

#### Library enrichment

15 µl of tagmented cDNA were mixed with 5 µl of 10 µM barcoded P7 primer (sequences provided in **Supplementary Table 1**). Per reaction, a mix of 28.5 µl nuclease-free water, 50 µl NEBNext High-Fidelity 2x PCR Master Mix (NEB #M0541S), 1 µl of 100x SYBR Green in DMSO (Life Technologies #S7563), and 0.5 µl of 100 µM Partial-P5 primer were added. Reactions were incubated in a qPCR cycler as follows: 72 °C for 3 min (for end fill-in after tagmentation), 98 °C for 30 s, cycle [98 °C for 10 s, 65 °C for 30 s, 72 °C for 30 s, plate read]. We monitored the fluorescence signals and removed samples from the thermocycler when they reached >4000 RFU. To complete unfinished PCR products, we incubated for 5 min at 72 °C in another thermocycler. Libraries were cleaned with a 0.7x SPRI cleanup with AMPure XP beads. All wells with the same P7 barcode were pooled and subjected to a 0.8x SPRI cleanup, eluting in 0.2 bead volumes of EB buffer (Qiagen #19086). The final library concentration was measured with the Qubit High Sensitivity DNA assay (Thermo Fisher Scientific #Q32854), and 1.5 ng were run on a Bioanalyzer High Sensitivity DNA chip (Agilent #5067-4626 and 5067-4627).

#### Sequencing

Libraries were diluted to 2.0 nM with EB buffer (Qiagen #19086) containing 0.1% Tween-20 (Sigma #P7949-500ML) and sequenced on the Illumina NovaSeq 6000 platform with standard sequencing primers and a read structure of 21 bases (Read 1), 8 bases (Index 1, i7), 16 bases (Index 2, i5), 78 bases (Read 2). Depending on the scale of the experiment, NovaSeq 6000 SP (Illumina #20027464), S1 (Illumina #20012865), or S2 (Illumina #20012862) reagent kits were used.

### Estimation of expected barcode collisions in scifi-RNA-seq

To obtain an estimate of cell doublets per compartment (droplet and/or well), we used the birthday problem as described by others^4, 20^, where the probability of collision of *n* cells in *c* compartments is defined as:

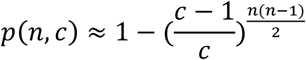

and the amount of cells per compartment *c* at a collision rate *p* is given by the reverse problem:

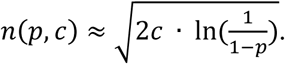

In this formulation, plate-based pre-indexing combined with subsequent barcoding of the same cells using a microfluidic droplet generator is equivalent to placing cells in a number of compartments corresponding to the product of the number of pre-indexing compartments and the number of droplets with beads in the microfluidic device. These estimates provide an informative upper bound for the number of usable single-cell transcriptomes, but are likely an overestimate due to higher-order collisions expected when loading high numbers of cells or nuclei.

### Monte Carlo simulation of microfluidic device loading

As a complementary approach that makes no assumptions about specific statistical distributions in the scifi-RNA-seq protocol, we performed the following Monte Carlo simulation: We generated two vectors of size *n* for which we randomly sampled integers from the range of pre-indexing compartments *r1* and microfluidic barcodes *r2*. We then calculated the fraction of multiple occurrences of the same *r1* integer in the *r2* compartments. This was repeated for a range of input cell numbers (*n*), compartments (*r1* and *r2*), and iterations (*i* = 100).

### Modeling of microfluidic device loading

The combinatorial estimation of expected barcode collisions as well as the Monte Carlo simulations do not adequately consider biological doublets (such as cell clumps) and should be regarded as upper bounds. To gain empirical insights into the loading of the Chromium system, we loaded nuclei without lysis reagents, such that they could be visualized inside the emulsion droplets under a light microscope (as described above). As we observed a fraction of droplets containing no nuclei even at high nuclei loading concentrations, we specified a Bayesian model where the observed number of nuclei per droplet is explained by a zero-inflated Poisson distribution of latent parameters *λ* for mean and variance and *Ψ* for the zero-inflated component. We imposed a prior on *λ* as the observed mean nuclei per droplet and estimated a posterior distribution of parameters by Markov Chain Monte Carlo (MCMC) sampling with the No U-Turn (NUTS) sampler for 200,000 initializer iterations, 4,000 tuning draws, and 4,000 draws to estimate the posterior parameters using PyMC3 (version 3.8)^30^.

To estimate doublet rates for scifi-RNA-seq experiments, we assumed that the final doublet rate is independent of the pre-indexing. In this case, for the purpose of doublet estimation, the number of nuclei that can produce doublets is the number of input nuclei divided by the number of unique pre-indexing barcodes. We used the model parameters estimated from the optically counted numbers of nuclei per droplet to estimate and interpolate model parameters across a range of nuclei loading concentrations. Using those parameters for realistic numbers of input nuclei, the doublet rate is estimated as one minus the probability mass function (PMF) of the zero-inflated Poisson distribution at a given nuclei loading concentration. While these estimates become reasonably accurate when actual nuclei counts observed for the microfluidic device are taken into account, the estimates are still somewhat optimistic as they are based on nuclei as the countable unit, whereas in the real scifi-RNA-seq experiment the countable units are transcript molecules, and the final number of nuclei are inferred from the transcript counting procedure.

### Processing of scifi-RNA-seq data

For fast parallel processing of raw sequence data, we demultiplexed the raw sequence calls based on the round1 barcode (and optionally a sample barcode if multiple scifi-RNA-seq experiments are multiplexed for sequencing), allowing up to 3 mismatches jointly for both barcode sequences; and the demultiplexed reads were written into unaligned single-read BAM files, with dedicated tags for the round2 barcode (“r2” tag) and UMI (“RX” tag).

The STAR aligner (version 2.7.0e) was used to map the demultiplexed unaligned BAM files with the following parameters: “-readFilesCommand samtools view -h -readFilesType SAM SE”. We allowed trimming of poly-A stretches by setting the “-clip3pAdapterSeq AAAAAA” parameter, and we retained all alignments via the following parameters: “-outFilterScoreMinOverLread 0 -outFilterMatchNminOverLread 0 -outFilterMatchNmin 0”. We set the output of STAR to aligned BAM using the “-outSAMtype BAM Unsorted” parameter.

To quantify gene expression, we used the featureCounts tool from the Subread package (version v1.6.2) to add an “XT” tag to the aligned BAM file with the respective gene using the “-g gene_id” parameter. Only alignments with quality above 30 were tagged (“-q 30” parameter). We quantified alignments in an unstranded manner (“-s 0” parameter) for either exons only (“-t exon” parameter) or for whole gene bodies (“-t gene” parameter) since nuclei can contain high levels of unspliced transcripts. We then used Pysam (version 0.15.2) to filter out alignments from low-quality or unmapped reads, from multi-mappers, and from secondary or supplementary alignments. Effectively, we kept only reads which aligned uniquely to genes with an alignment quality score above 30.

To generate a gene expression matrix, we considered a unique molecule count as a sequenced molecule with unique round1 and round2 barcodes, UMI, associated gene, and mapping position in the chromosome. We then created a Scipy (version 1.3.0) compressed sparse row matrix of the form (UMI count, cell barcode, gene) and initialized an Anndata (version 0.6.21) object to save it as h5ad format. We used Python 3.7.2 for all analysis.

### Analysis of scifi-RNA-seq data

We used Scanpy (version 1.4.3) for further analysis of the single-cell RNA-seq data. For the cell line mixture and T-cell experiments, we filtered the Anndata object for cell barcodes containing at least 200 UMI counts, while keeping only those genes that were not in the lower 10% percentile of cumulative UMI counts and that were detected by least 500 barcodes. We further excluded cells more than three standard deviations away from the mean of the fraction of the transcriptome in ribosomal or mitochondrial genes. Gene expression values were normalized per cell total in order to sum to 10,000, and log-transformed. We performed principal component analysis on scaled and centered expression values, computed a neighbor graph, and performed dimensionality reduction with UMAP using the respective scanpy functions, all with default parameters. We then employed the Leiden algorithm for community detection on the neighbor graph with the 0.5 resolution parameter, and we tested for mean gene expression differences between groups of cells using the normalized and log transformed expression values and the *scanpy*.*tl*.*rank_genes_groups* function with a two-sided t-test with variance overestimation (*t-test_overstim_var* function) and corrected for multiple testing with the Benjamini-Hochberg procedure. For these differential gene sets between cell lines, we performed gene set enrichment analysis using the Enrichr API.

### Data availability

All data will be available through the Supplementary Website (http://scifi-rna-seq.computational-epigenetics.org), which is currently in preparation. Moreover, the single-cell RNA-seq data will be available from the NCBI GEO database.

### Code availability

The analysis source code underlying the final version of the paper will be available on the Supplementary Website (http://scifi-rna-seq.computational-epigenetics.org) and in a Git repository (https://github.com/epigen/scifiRNA-seq).

## Acknowledgements

We thank Martin Senekowitsch and the team of the Biomedical Sequencing Facility at CeMM for assistance with next generation sequencing and all members of the Bock lab for their help and advice. C.B. is supported by a New Frontiers Group award of the Austrian Academy of Sciences and by an ERC Starting Grant (European Union’s Horizon 2020 research and innovation programme, grant agreement n° 679146).

## Author contributions

P.D. conceived and developed the method; P.D. and T.B. optimized the protocol and conducted the experiments; A.F.R performed computational modeling and analyzed the data; P.D. and A.F.R visualized the results and prepared the figures; T.K. contributed to the T cell experiments; D.B. performed data pre-processing; P.D., A.F.R., and C.B. wrote the manuscript with contributions from all authors; C.B. supervised the project.

## Competing financial interests

P.D. and C.B. are inventors on a patent application describing scifi barcoding and the scifi-RNA-seq method.

## Supplementary figures

**Supplementary Figure 1:**
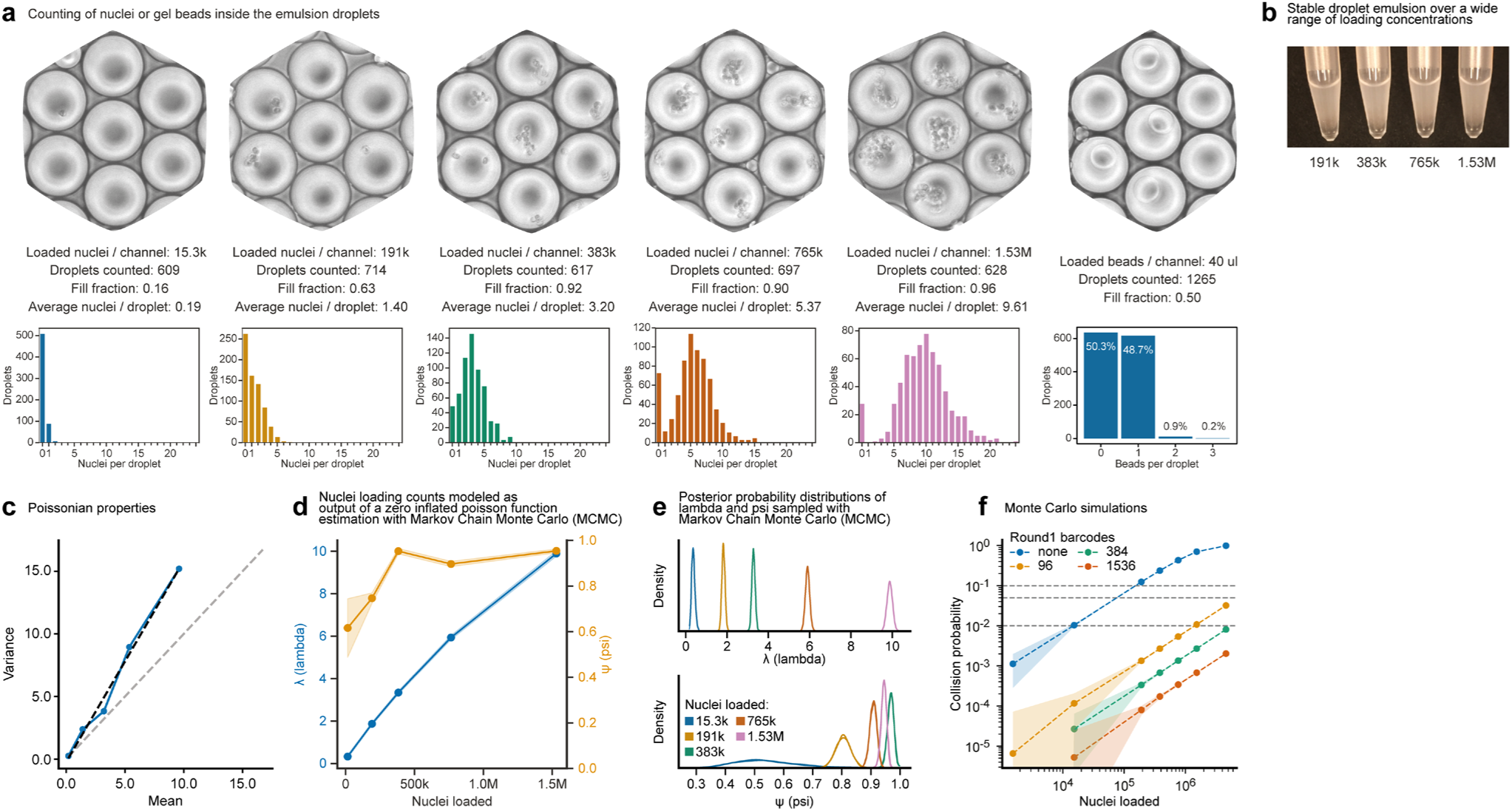
Droplet overloading on the Chromium microfluidic droplet generator. **a)** By omitting the lysis reagents, nuclei remained intact and were imaged using a standard microscope, allowing the counting of nuclei per droplet. Results for loading concentrations of 15,300, 191,000, 383,000, 765,000, and 1,530,000 nuclei per channel are summarized as histograms. For each loading concentration, the number of evaluated droplet images, the droplet fill fraction, and the average number of nuclei per droplet are shown. Furthermore, by substituting the nuclei suspension with 1x Nuclei Buffer and omitting Reducing Agent B, intact gel beads were visualized inside the emulsion droplets. Bead fill rates based on 1,265 evaluated droplet images are shown. **b)** Despite substantial droplet overloading, we obtained stable droplet emulsions for all tested conditions. **c)** Nuclei loading displays properties of a Poisson-like distribution. **d)** Computational modeling of nuclei loading as a zero-inflated Poisson function. **e)** Posterior probability distributions of lambda and psi sampled with Markov Chain Monte Carlo (MCMC). **f)** Independent estimation of the cell doublet rates through Monte Carlo simulations.

**Supplementary Figure 2:**
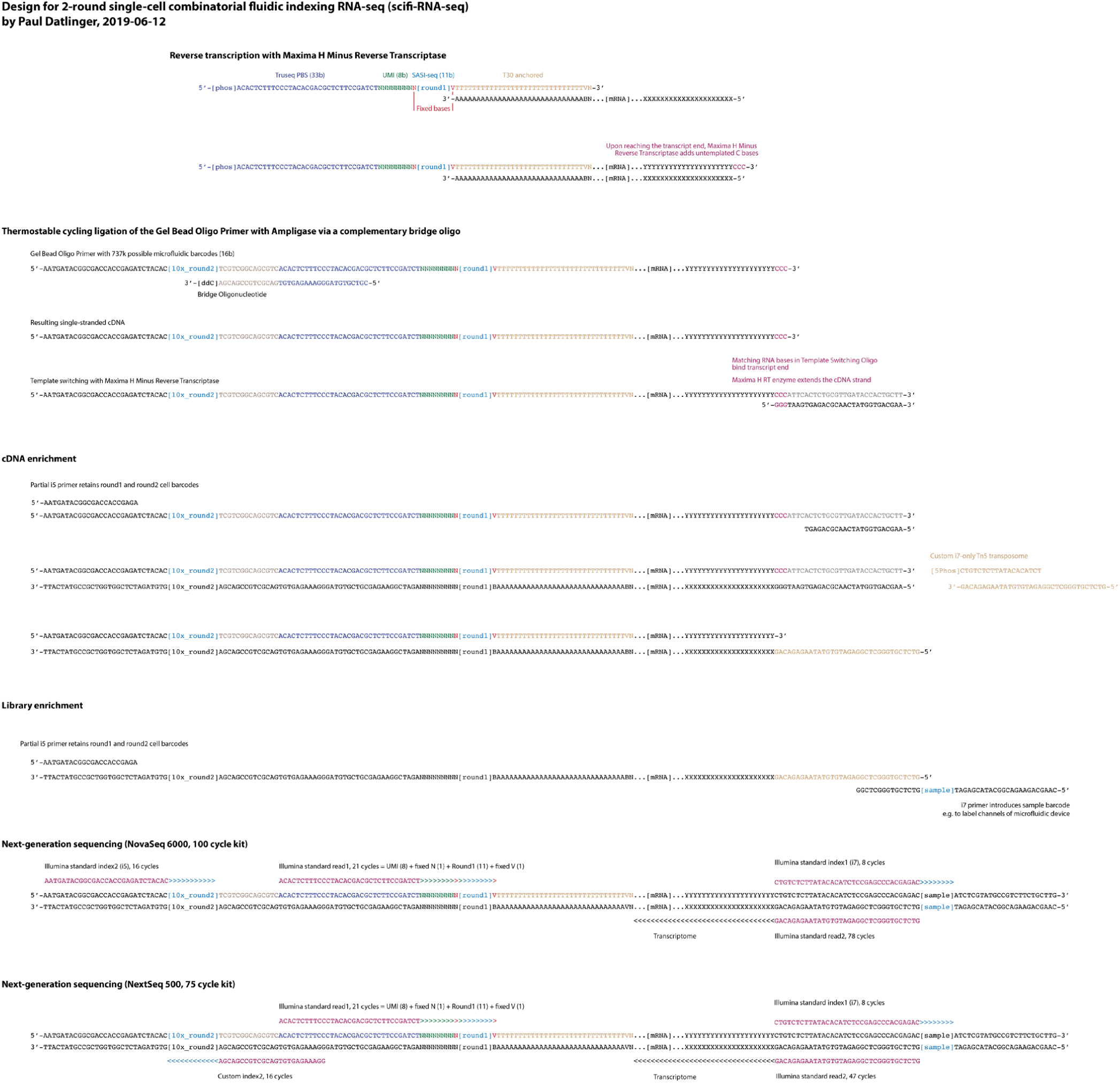
Detailed method design for scifi-RNA-seq including sequence information. Schematic outline of scifi-RNA-seq with detailed oligonucleotide sequences. The reverse transcription is performed inside intact cells or nuclei on a microwell plate, introducing well-specific round1 barcodes into the whole transcriptome. Pre-indexed cells or nuclei are pooled and encapsulated into emulsion droplets using a standard microfluidic droplet generator. The round2 barcodes are introduced by thermocycling ligation with a complementary bridge oligo and thermostable ligase. Afterward, the droplet emulsion is broken, and a second defined end is introduced into the library via template switching. cDNA is enriched and tagmented with a custom i7-only transposome. Finally, the library is PCR-enriched, with the option to introduce an additional sample index. The read structure for next-generation sequencing on the Illumina NovaSeq 6000 and NextSeq 500 platforms is shown.

**Supplementary Figure 3:**
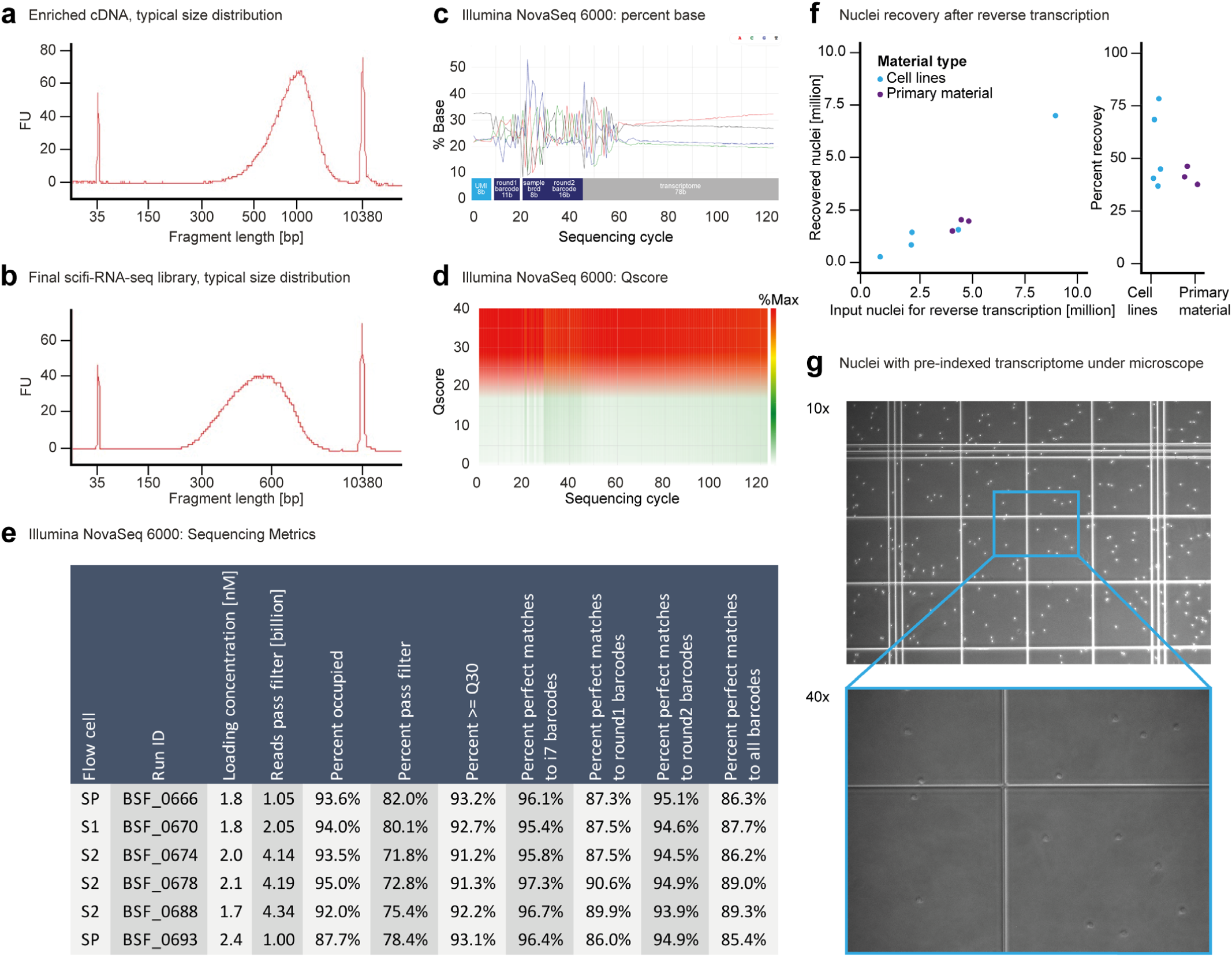
scifi-RNA-seq library preparation and next-generation sequencing. **a)** Typical size distribution of enriched cDNA obtained using scifi-RNA-seq. **b)** Typical size distribution of a final scifi-RNA-seq library ready for next-generation sequencing. **c)** Distribution of DNA bases along scifi-RNA-seq sequencing reads, showing the characteristic sequence patterns of the UMI, round1 barcode, round2 barcode, sample barcode, and transcript. **d)** Heatmap showing sequencing quality (Qscore) for each sequencing cycle. **e)** Table summarizing all NovaSeq 6000 sequencing runs performed as part of this study. scifi-RNA-seq was thoroughly tested with NovaSeq SP, S1, and S2 reagents. The table also summarizes the percentage of reads with perfect match to the sample (i7) barcode, pre-indexing (round1) barcode, microfluidic (round2) barcode, and with a correct combination of all three barcodes. **f)** Nuclei recovery after pre-indexing of the whole transcriptome by reverse transcription. scifi-RNA-seq achieves high recovery rates for both cell lines and primary material. **g)** Nuclei with pre-indexed transcriptome, prior to microfluidic device loading, visualized under a microscope in a counting chamber. The selected images show nuclei derived from human primary T cells.

**Supplementary Figure 4:**
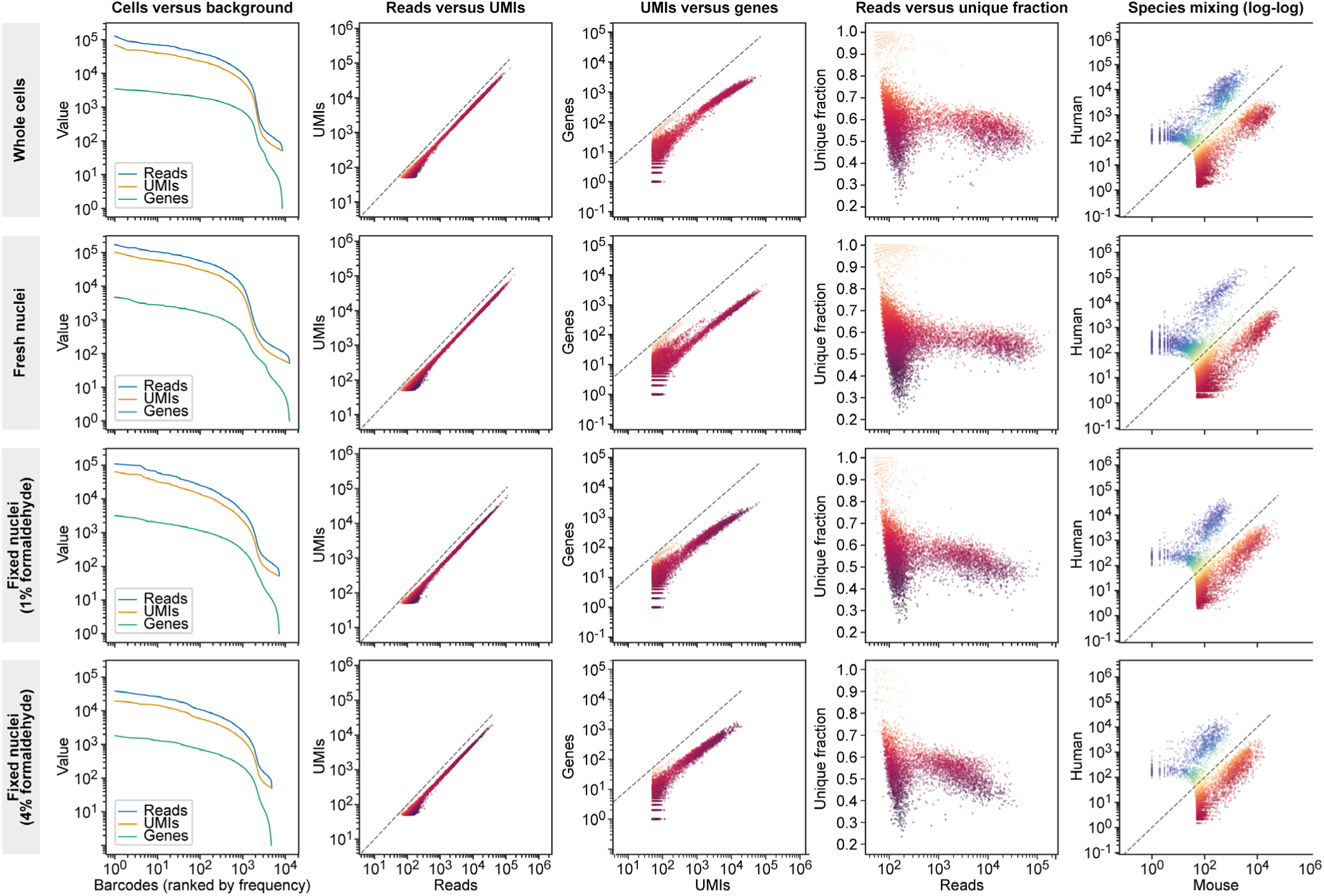
scifi-RNA-seq yields high-quality data for whole cells, fresh nuclei, and fixed nuclei. We prepared a mixture of human (Jurkat) and mouse (3T3) cells, and performed scifi-RNA-seq on whole cells permeabilized by methanol, freshly isolated nuclei, and nuclei fixed with 1% or 4% formaldehyde that were cryopreserved, re-hydrated, and permeabilized. During reverse transcription on a 96-well plate, each sample was assigned a specific set of round1 barcodes. Afterward, all wells were pooled, and 15,300 cells/nuclei were loaded into a single channel of the Chromium device. We provide the following performance plots: (i) ranked barcodes plotted against reads, unique molecular identifiers (UMIs), or detected genes, distinguishing single-cell transcriptomes from background noise; (ii) reads plotted against UMIs; (iii) reads plotted against the number of detected genes; (iv) reads plotted against the fraction of unique reads; (v) species mixing plot showing the number of UMIs per cell aligning to the mouse genome (x-axis) versus the human genome (y-axis). To facilitate comparisons between different types of input material, the axes of the performance plots use the same scale across conditions.

**Supplementary Figure 5:**
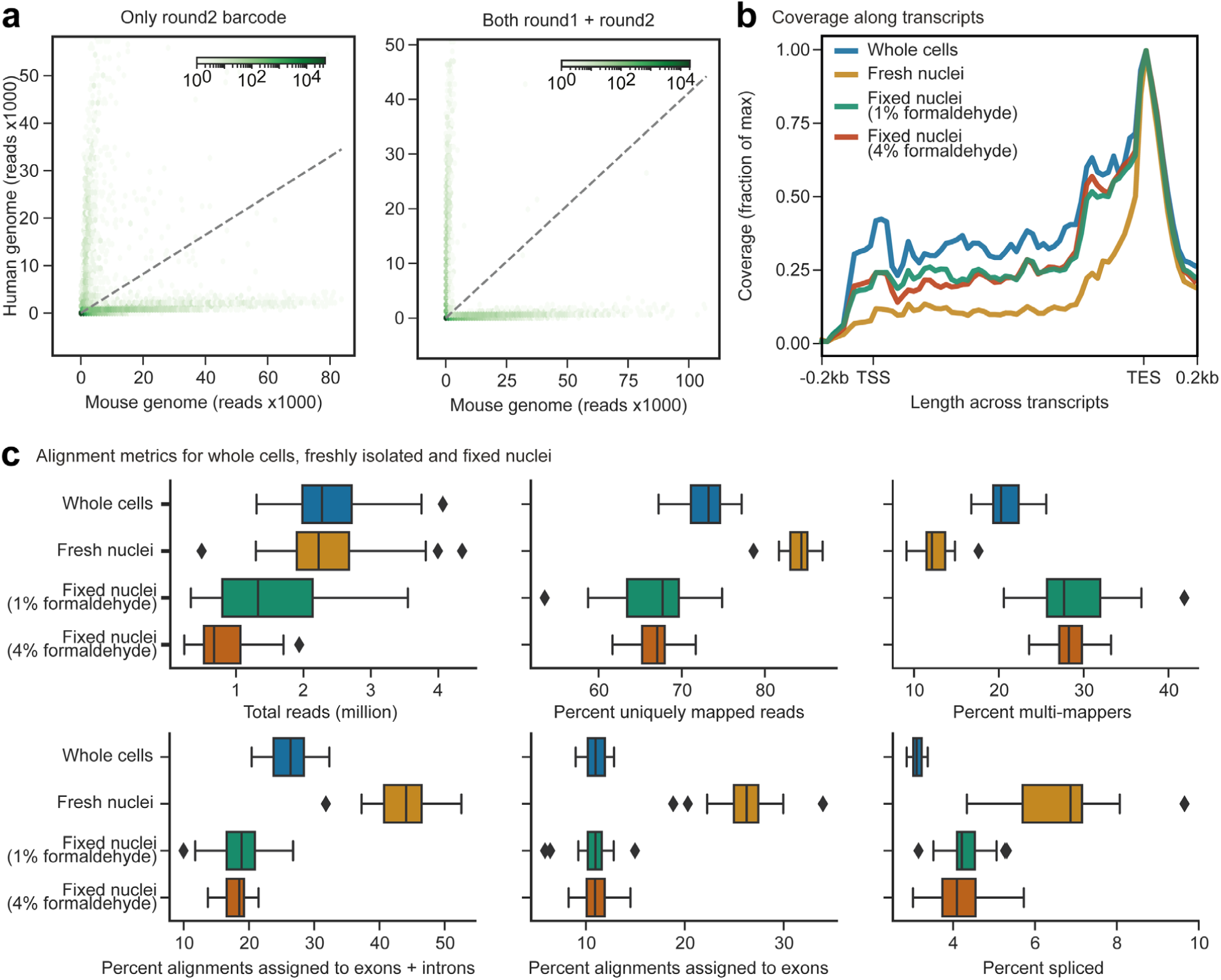
scifi-RNA-seq alignment metrics for whole cells, fresh nuclei, and fixed nuclei. **a)** 15,300 pre-indexed nuclei from a mixture of human (Jurkat) and mouse (3T3) cells were processed in a single microfluidic channel and demultiplexed based on the microfluidic round2 barcode only (left plot), or based on the combination of round1 and round2 barcodes (right plot). At the standard loading concentration of the Chromium device (15,300 nuclei per channel), the microfluidic (round2) index provides sufficient complexity to resolve single cells, although the combination of round1 and round2 barcodes still results in a reduction of background noise. **b)** Coverage along human and mouse transcripts from 200 bp upstream of the transcription start site (TSS) to 200 bp downstream of the transcription end site (TES), shown for whole cells permeabilized by methanol, freshly isolated nuclei, and nuclei fixed with 1% or 4% formaldehyde that were cryopreserved, re-hydrated, and permeabilized. Freshly isolated nuclei show the strongest 3’ enrichment. **c)** Boxplots summarizing sequence alignment metrics across the different types of input material: Total reads sequenced, percent uniquely mapped reads, percent multi-mappers, percent alignments to exons plus introns, percent alignments to exons, and percent spliced reads. Freshly isolated nuclei showed the best performance for these alignment metrics.

**Supplementary Figure 6:**
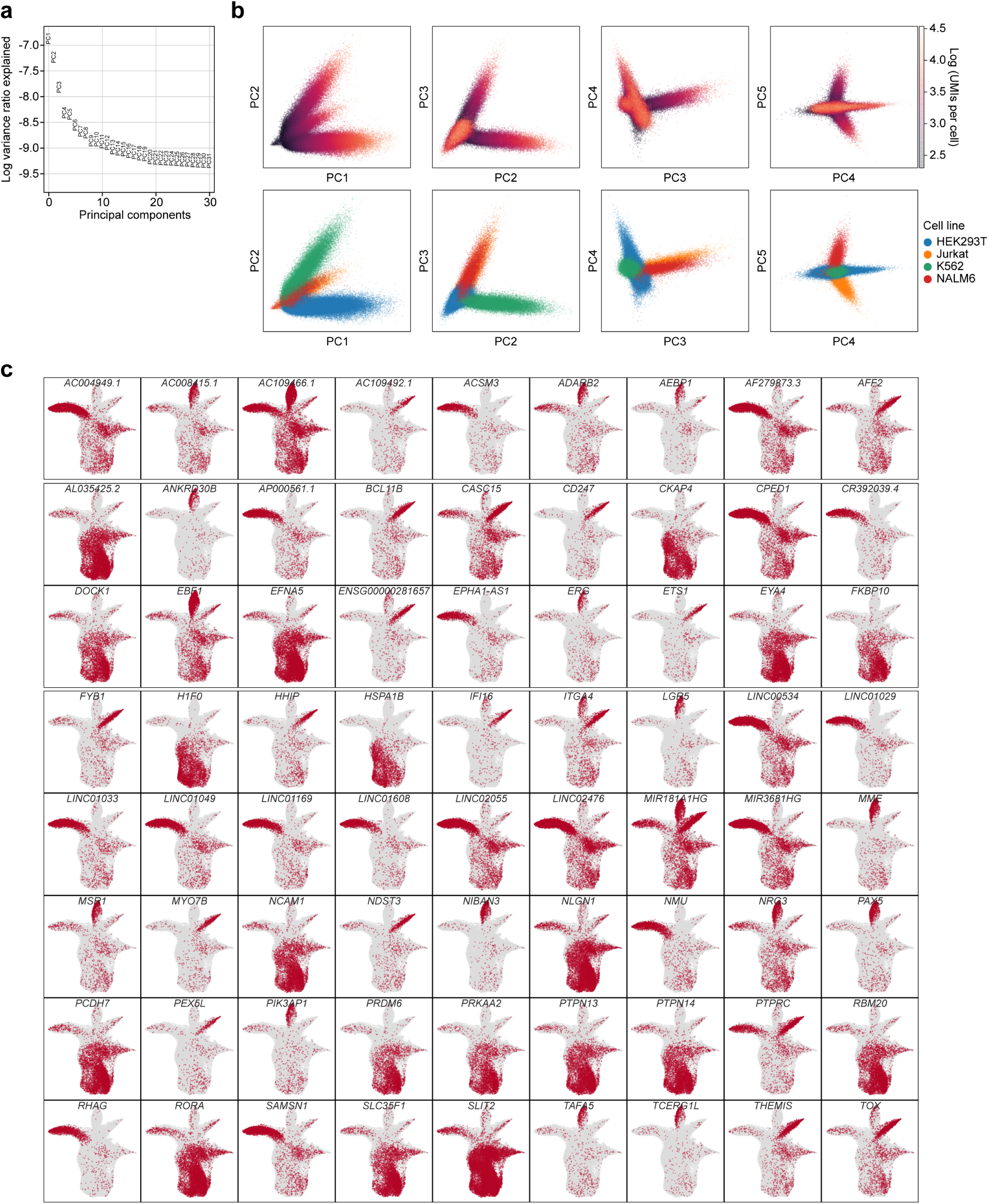
scifi-RNA-seq profiling for a mixture of four human cell lines. **a)** Variance explained by the top 30 principal components. **b)** Principal component analysis (PCA) projections for 151,788 single cells, color-coded with the number of UMIs per cell (top row) and with round1 barcodes denoting cell lines. **c)** Expression values of 72 additional cell line specific genes mapped onto the UMAP projection as shown in Fig. 2f.

**Supplementary Figure 7:**
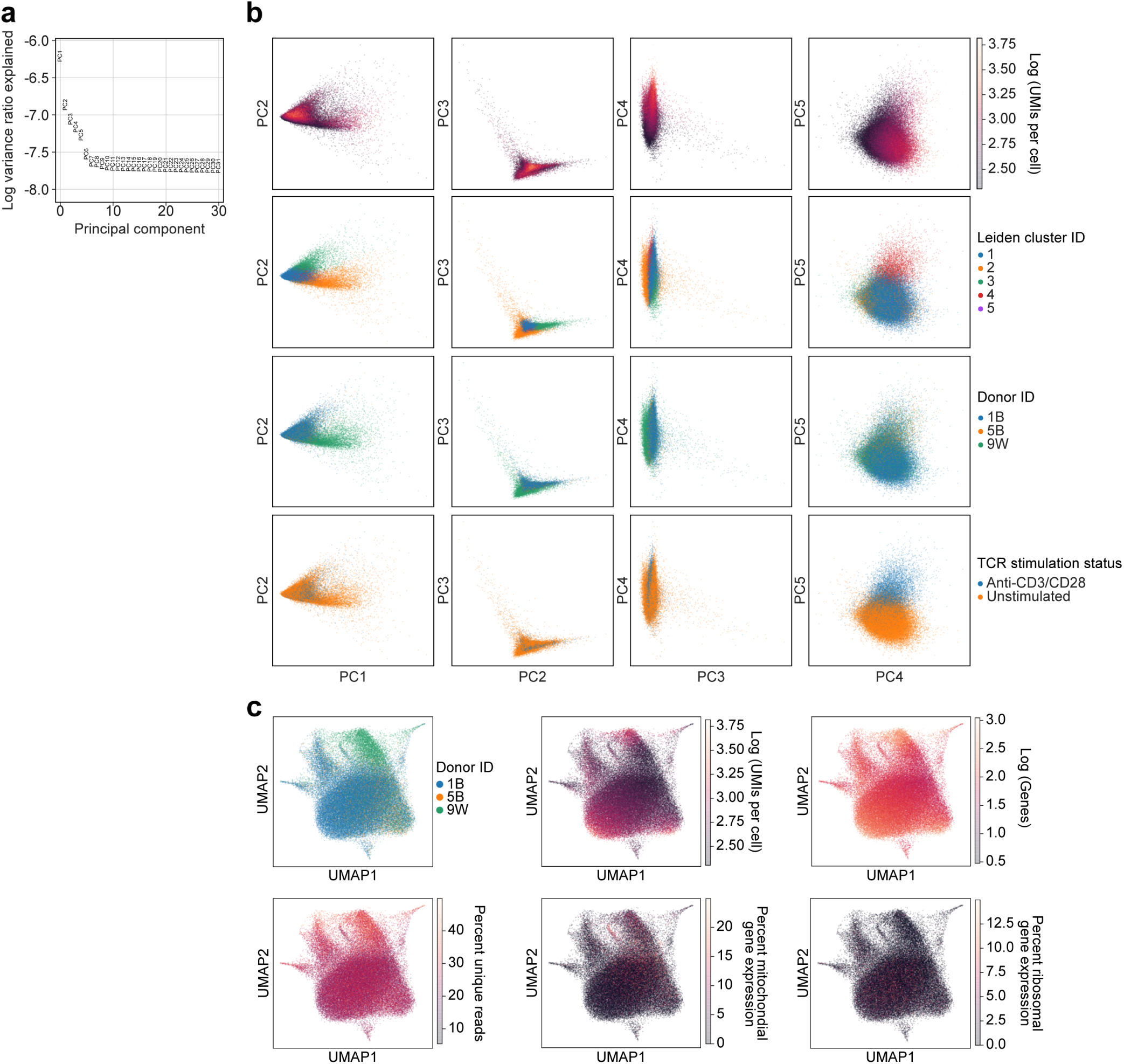
scifi-RNA-seq profiling for primary T cells with and without TCR stimulation. **a)** Variance explained by the top 30 principal components. **b)** PCA projections for 62,558 single cells. From top to bottom, the following variables are mapped onto these projections: Logarithm of UMIs per cell, cluster ID, donor ID, and T cell receptor (TCR) stimulation status. **c)** UMAP projections for 62,558 single cells (as shown in Fig. 2i) with additional variables mapped onto the projections: Donor ID, logarithm of UMIs per cell, logarithm of detected genes per cell, percent unique reads per cell, percent mitochondrial expression, and percent ribosomal expression.

